# Synergistic epistasis enhances cooperativity of mutualistic interspecies interactions

**DOI:** 10.1101/2020.06.22.160184

**Authors:** Serdar Turkarslan, Nejc Stopnisek, Anne W Thompson, Christina E Arens, Jacob J Valenzuela, James Wilson, Kristopher A Hunt, Jessica Hardwicke, Sujung Lim, Yee Mey Seah, Ying Fu, Liyou Wu, Jizhong Zhou, Kristina L Hillesland, David A Stahl, Nitin S Baliga

## Abstract

Frequent fluctuations in sulfate availability rendered syntrophic interactions between the sulfate reducing bacterium *Desulfovibrio vulgaris* (*Dv*) and the methanogenic archaeon *Methanococcus maripaludis* (*Mm*) unsustainable. By contrast, prolonged laboratory evolution in obligate syntrophy conditions improved the productivity of this community but at the expense of erosion of sulfate respiration (SR). Hence, we sought to understand the evolutionary trajectories that could both increase the productivity of syntrophic interactions and sustain SR. We combined a temporal and combinatorial survey of mutations accumulated over 1000 generations of 9 independently-evolved communities with analysis of the genotypic structure for one community down to the single-cell level. We discovered a high level of parallelism across communities despite considerable variance in their evolutionary trajectories and the perseverance of a rare SR+ *Dv* lineage within many evolution lines. An in-depth investigation revealed that synergistic epistasis across *Dv* and *Mm* genotypes had enhanced cooperativity within SR- and SR+ assemblages, allowing their co-existence as *r*- and *K*-strategists, respectively.

## INTRODUCTION

Syntrophic interactions between bacteria and archaea are a major driver of anaerobic transformations of >1 gigaton/yr of C into methane, which is ∼30 times more potent than CO_2_ as a greenhouse gas, and also a sustainable fuel (Thauer, 2011). Across diverse anoxic environments, including anaerobic reactors, animal guts, ocean and lake sediments and soils, in the absence of respirable electron acceptors such as nitrate and sulfate, diverse syntrophs partner with methanogens to oxidize organic material. Syntrophy lifestyle can be either obligate or facultative, for example, although while oxidation via sulfate respiration (SR) is energetically favorable compared to syntrophy, many Sulfate Reducing Bacteria (SRBs) are facultative syntrophs that conditionally engage in syntrophy with methanogens in the absence of sulfate (Oude Elferink et al., 1994).

Conditional switching between syntrophy and SR is energetically expensive requiring the differential regulation of thousands of genes. Not surprisingly, frequent fluctuations between SR and syntrophy was demonstrated to be energetically unsustainable for a coculture of *Desulfovibrio vulgaris* Hildenborough (*Dv*) and *Methanococcus maripaludis* S2 (*Mm*) causing the syntrophic community to collapse with as few as 4 transitions (Turkarslan et al., 2017). By contrast, prolonged laboratory evolution of the same community under obligate syntrophy conditions resulted in significantly improved growth and stability within 300 generations, but at the expense of loss of independence through the erosion of SR (Hillesland and Stahl, 2010, Hillesland et al., 2014). While SR eroded across nearly all evolution lines, other processes such as regulation and signal transduction also accumulated mutations suggesting that modulation of multiple pathways could have also contributed to improved syntrophic growth. This was not surprising because syntrophic mutualism is known to employ many processes including diffusion of shared metabolites (Rotaru et al., 2012), interspecies electron transfers (McGlynn et al., 2015) and aggregation of cells for efficient cross-feeding (Summers et al., 2010).

Another striking discovery was that a subpopulation of cells capable of respiring sulfate (SR+) persisted in low frequency within the dominant non-sulfate respiring (SR-) populations for most evolved lines (Hillesland et al., 2014). Persistence of SR+ cells during syntrophy suggested that they may be adapted to a narrow niche that the dominant SR-population is unable to exploit effectively. One hypothesis is that SR+ and SR-have specialized on growth dynamics, allowing for their co-existence, e.g., as *r*- and *K*-strategists in a seasonal environment (Wei and Zhang, 2019, Pianka, 1970, Rozen and Lenski, 2000). Another consideration is that the Black Queen Hypothesis (BQH) can explain the persistence of the SR+ population (Morris et al., 2012). In this hypothesis the SR+ population subsists by producing a costly metabolite that SR-cells cannot. Dependency of SR-cells on the metabolite prevents them from completely excluding SR+ cells even though the SR+ cells pay the cost for the metabolite. Given that mutations in many pathways in the two organisms could have improved the mutualism, this also raised the possibility that distinct cooperative interactions between SR- and SR+ populations and different subpopulations of evolved *Mm* (partner choice (Archetti, 2011)) could have independently increased productivity of syntrophy in each of the two subpopulations. Notably, naturally occurring polymorphisms in the ion-translocating subunit cooK of membrane-bound COO hydrogenase of *Dv* are known to be essential for mutualism with *Mm*, demonstrating that partner choice is important in promoting facultative syntrophic interactions (Großkopf et al., 2016). The *Dv* and *Mm* syntrophic community therefore offers a unique opportunity to elucidate and characterize the evolutionary trajectories and mechanisms that increase the productivity of mutualistic interactions among microbes that co-exist in diverse environments (e.g., gut, soil, etc.) and play a central role in an important step in biogeochemical C cycling.

Insights into evolutionary trajectories for increased productivity of mutualistic interactions have primarily been elucidated using synthetic communities built from laboratory constructed auxotrophic strains of *Escherichia coli* (Mee et al., 2014), or yeast (Shou et al., 2007), and cocultures of *E. coli* and *Salmonella typhimurium* (Douglas et al., 2017). In addition to discovery of numerous evolutionary phenomena for improvement of mutualistic interactions, these studies have demonstrated the potential for the emergence of synergistic epistasis and cooperation within microbial communities. Here, we have investigated whether synergistic epistasis and cooperativity can emerge through the selection of interactions among specific genotypes during the evolution of environmentally-important syntrophy, wherein one partner (*Dv*) generates a product (H_2_) that inhibits its own growth, and the second partner (*Mm*) consumes the byproduct to promote growth of both organisms.

Briefly, building on our prior work, in this study we tracked the sequence and combinations in which mutations accumulated in *Dv* and *Mm* across 100, 300, 500, 780 and 1K generations across nine independent evolution lines (five cultured with shaking and four without). From the 1K generation of three lines, we generated simplified communities through serial end-point dilutions (EPDs). All simplified communities demonstrated growth characteristics comparable to their more complex parental communities, suggesting that evolved variants and interactions essential for the community phenotype were retained. Bulk sequencing of the simplified communities was then used to characterize how parental mutations were segregated into each EPD. Together, the combinations and temporal distributions of mutations across generations and EPDs discovered evidence for the existence of interactions among specific evolved lineages of *Dv* and *Mm*, within the same evolution line. Through single cell sequencing, we then inferred and characterized interactions within a SR+ and a SR-EPD derived from the same parental population. Finally, we quantified growth characteristics (growth rate, yield, and cooperativity) of each EPD and pairings of evolved clonal isolates of *Dv* and *Mm* with each other and the ancestral strains. These analyses uncovered synergistic epistasis as a plausible mechanism for the increased cooperativity of mutualistic interactions within EPDs, explaining how SR+ and SR-subpopulations co-exist as *r*- and *K*-strategists (**Fig 1**).

**Figure 1.**
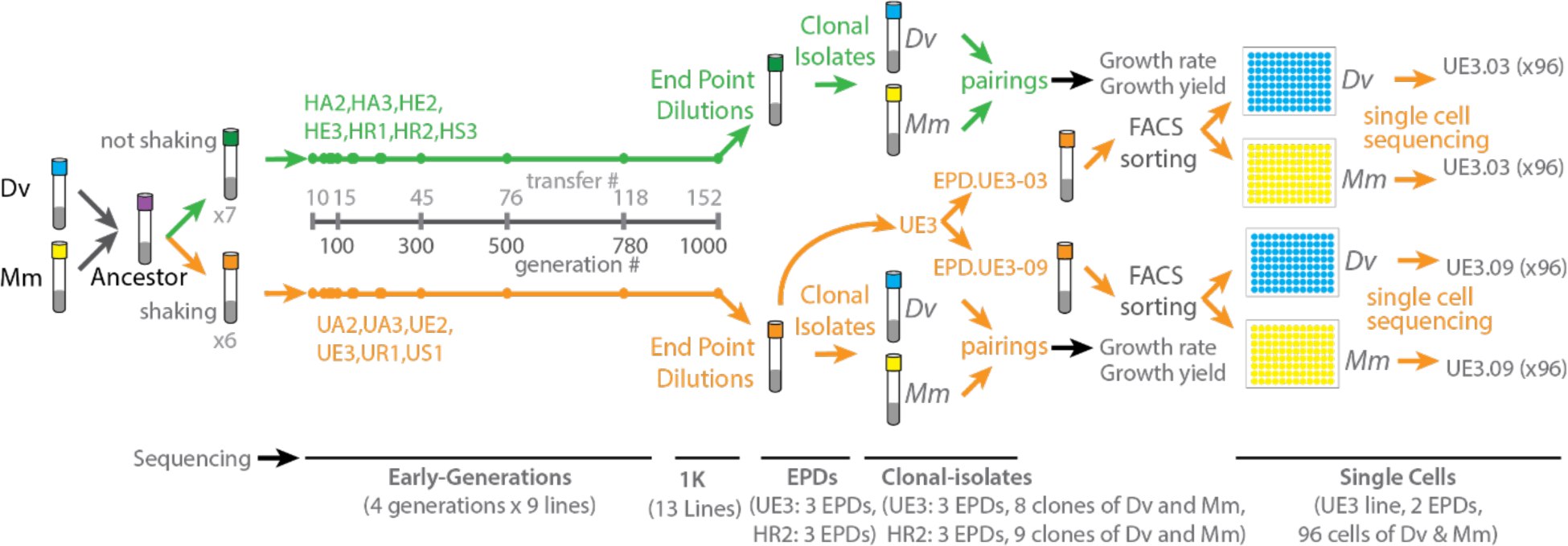
Overview of directed laboratory evolution to probe evolutionary signatures for syntrophic cocultures of *Dv* and *Mm*. Thirteen independent cocultures were subjected to laboratory evolution with and without shaking as described before (Hillesland et al., 2014). DNA samples were collected across generations, End-Point-Dilutions (EPDs), clonal isolates, and single cells to identify genomic alterations. In addition, clonal isolates were paired in varied combinations in order to determine growth rate and yield for cocultures. Number of samples sequenced are indicated at the bottom.

## RESULTS

### Distribution, frequency, and functional implications of mutations during laboratory evolution of obligate syntrophy

We evaluated whether selection of mutations in the same genes (i.e., “parallel evolution” (Stern, 2013)) had contributed to improvements in syntrophic growth of *Dv* and *Mm* across independent evolution lines, all of which started with the same ancestral clone of each organism. Based on the frequency of mutations (normalized to gene length and genome size) in *Dv* and *Mm* across 13 evolved lines (six lines designated U for “uniform” conditions with continuous shaking and seven H lines for “heterogenous” conditions without shaking), we calculated a G-score (Tenaillon et al., 2016) (“goodness-of-fit”, see Methods (Tenaillon et al., 2016)) to assess if the observed parallel evolution rate was higher than expected by chance. The “observed G-score” was calculated as the sum of G-scores for all genes in the genome of each organism; mean and standard deviation of “expected G-scores” were calculated through 1000 simulations of randomizing locations of observed numbers of mutations across the genome of each organism. The observed total G-score for *Dv* (1092.617) and *Mm* (805.02) was significantly larger than the expected mean G-score (*Dv*: 798.19 ± 14.99, Z=19.63 and *Mm*: 564.83 ± 15.95, Z=15.06), which supported significant convergence in genotypic evolution across laboratory evolution lines.

Altogether, 24 genes in *Dv* [G-score range: 7.7 (DVU1012) to 156.6 (DVU0799)] and 16 genes in *Mm* [range: 10.1 (MMP1363) to 166.4 (MMP1718)] had accumulated function modulating mutations across at least 2 or more evolution lines (**Fig 2**). Notably, mutations within the same gene were in different locations and they appeared at different times across independent lines, further supporting parallel evolution. The 40 genes implicated in parallel evolution represent core functions, including signal transduction and regulation (7 in *Dv* and 6 in *Mm*), SR (4 in *Dv*), transport (4 in *Dv* and 3 in *Mm*), and motility (1 each in *Dv* and *Mm*) (Tenaillon et al., 2012, Kvitek and Sherlock, 2013). Mutations in SR genes were among the top contributors to the total G-score in *Dv* (DVU2776 (74.7), DVU1295 (46.5), DVU0846 (42.9), and DVU0847 (22.3)). We had demonstrated previously that obligate requirement of mutual interdependence drove the erosion of metabolic independence of *Dv* through accumulation of SR-mutations (Hillesland et al., 2014), a well-known phenomenon that evolution in a uniform and limited resource environment selects for specialists (Van den Bergh et al., 2018). However, it was intriguing that DVU2776 (DsrC), which catalyzes conversion of sulfite to sulfide, the final step in SR, accumulated function modulating mutations across 6 lines, suggesting that these changes might promote some alternate function for this protein, including electron confurcation for the oxidation of lactate (Meyer et al., 2013), sulfite reduction, 2-thiouridine biosynthesis and possibly gene regulation (Venceslau et al., 2014). Notably, we have demonstrated previously that while SR was universally lost across all lines, the SR-mutants could be recovered on lactate-sulfite agar plates (Hillesland et al., 2014).

**Figure 2:**
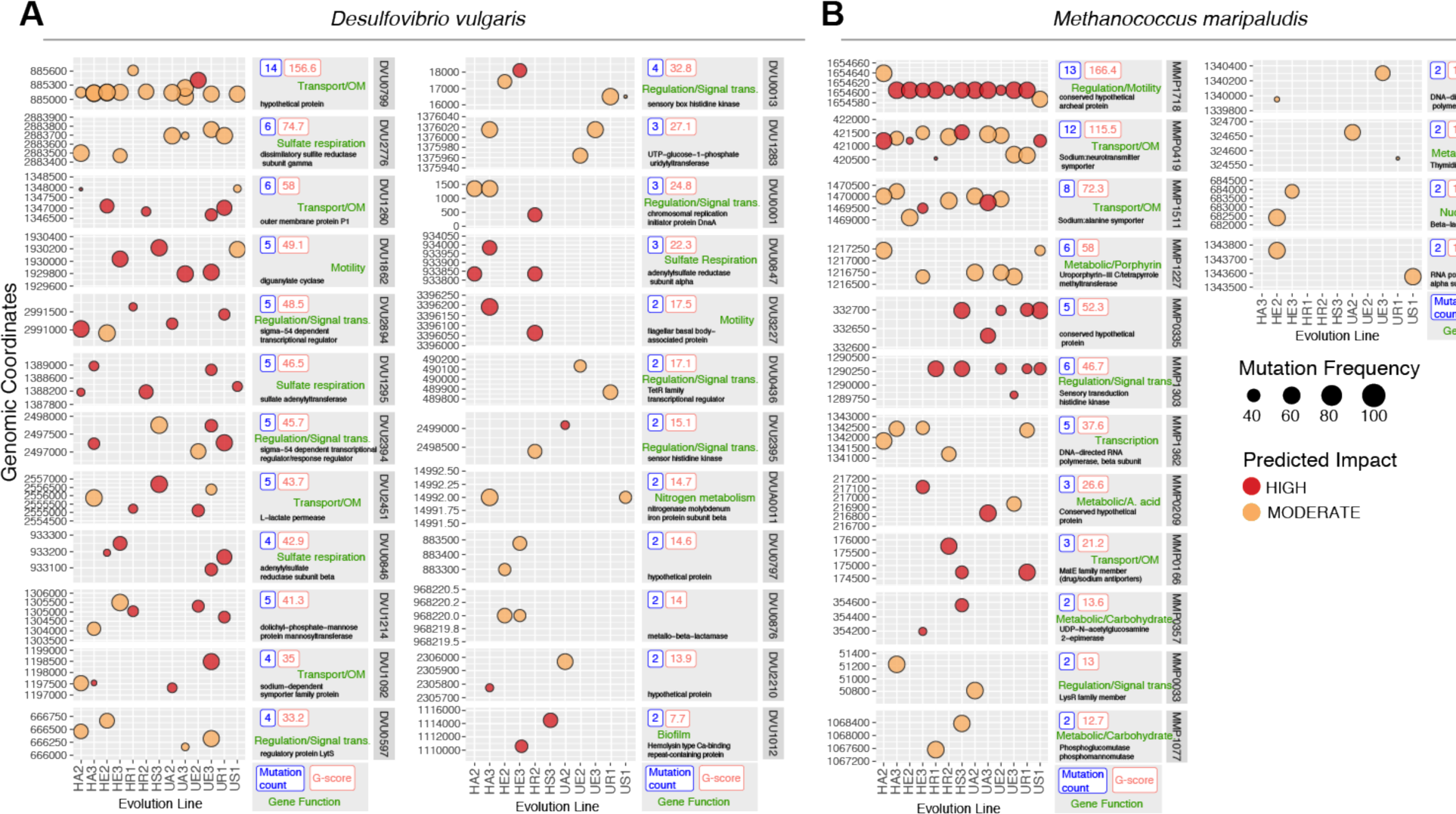
Frequency and location of high G-score mutations in *Dv* and *Mm* across 13 independent evolution lines. SnpEff predicted impact of mutations* are indicated as moderate (orange circles) or high (red circles) with frequency of mutations indicated by node size. Expected number of mutations for each gene were calculated based on the gene length and the total number of mutations in a given evolution line. Genes with parallel changes were ranked by calculating a G (goodness of fit) score between observed and expected values and indicated inside each panel. Mutations for each gene are plotted along their genomic coordinates (vertical axes) across 13 evolution lines (horizontal axes). Total number of mutations for a given gene is shown as horizontal bar plots. [*HIGH impact mutations: gain or loss of start and stop codons and frame shift mutations; MODERATE impact mutations: codon deletion, non-synonymous in coding sequence, change or insertion of codon; low impact mutations: synonymous coding and non-synonymous start codon]

Signal transduction and regulatory gene mutations represented 19.9% and 27.2% of all mutations in *Dv* and *Mm*, respectively, echoing similar observations during long term laboratory evolution of *E. coli* (Tenaillon et al., 2016), potentially because of their influence on the functions of large numbers of genes (Barrick et al., 2009, Cooper et al., 2003). Not surprisingly, five of these genes (DVU0597, DVU1862, DVU0436, DVU0013, and DVU2394) also accumulated mutations during long term salt adaptation of *Dv*, suggesting that these adaptive changes were not specific to syntrophic interactions with *Mm*. Given that salinity of the syntrophic medium was higher than that routinely used for culturing *Dv*, these mutations are potentially relevant for adaptation to higher salt environments (Zhou et al., 2017, Zhou et al., 2015). DVU2394, for instance, is a member of a two component system (with DVU2395) for regulating energy metabolism genes (DVU2405-DVU2397) that might be important for salinity adaptation (Rajeev et al., 2011). On the other hand, 8 intergenic regions and 45 genes (18 high G-score genes) that accumulated mutations across different lines appeared to be specific to syntrophic interactions and of diverse functions ([SR: DVU0846, DVU0847, DVU1295, DVU2776], [transport/outer membrane: DVU0799, DVU2451, DVU1092], [motility: DVU3227], [metabolism: DVU1214, DVU1283, DVUA0011], [RNA-degradation: DVU0876], [regulation/signal transduction: DVU2894, DVU0001, DVU2395], [unknown function: DVU0797, DVU2210], [biofilm formation: DVU1012]).

Five evolution lines were cultured with continuous shaking (“uniform” conditions, U lines), whereas four were cultured without shaking (“heterogeneous” conditions, H lines) to investigate the role of motility and aggregation in promoting cross-feeding. Notably, the regulator of the archaellum operon (MMP1718) had the highest G-score with frameshift (11 lines) and nonsynonymous coding (2 lines) mutations (Ding et al., 2016). Similarly, two motility-associated genes of *Dv* (DVU1862 and DVU3227) also accumulated frameshift, nonsense and non-synonymous mutations across 4 H and 3 U lines. Together, these observations demonstrated that retaining motility might have had a fitness cost in the evolution of syntrophy across both H and U lines, consistent with the outcome of other laboratory evolution experiments where species were also propagated in liquid media (Velicer et al., 2002) and with estimates of motility costs under energy limitation (Martínez-García et al., 2014, Kempes et al., 2017).

Missense and nonsense mutations in outer membrane and transport functions (4 genes in *Dv* and 3 genes in *Mm*) might have also promoted cross-feeding. The highest G-score gene in *Dv*, DVU0799 –an abundant outer membrane porin for uptake of sulfate and other solutes in low sulfate conditions (Zeng et al., 2017), was mutated early across all lines, with at least two missense mutations in UE3 (S223Y) and UA3 (T242P) potentially disrupting predicted phosphorylation sites. While it was to be expected that *Dv*, a generalist, would shed several functions that were not relevant for syntrophy, it was surprising that there was further specialization of the *Mm* genome that is believed to have evolved in an anaerobic syntrophic environment. This additional genome streamlining could be a result of natural selection of complementary capabilities to improve metabolic interdependency and cross-feeding between *Dv* and *Mm* (Martinez-Cano et al., 2014, McNally and Borenstein, 2018). For instance, the loss of function and missense mutations in MMP1511, a Na/Ala symporter was consistent with its downregulation during syntrophy and its potential role in alanine metabolism mediated syntrophic coupling (Walker et al., 2012). Na symport of alanine would deplete the sodium motive force, so there could be an energetic advantage to mutations conferring passive uptake. In summary, parallel evolution in motility, regulation, signal transduction, and transport functions across both *Mm* and *Dv* likely contributed to improvement in growth characteristics during syntrophy.

### Analysis of temporal appearance and combinations of mutations across evolution lines

Growth characteristics of all evolution lines improved by the 300^th^ generation (Hillesland and Stahl, 2010), and in some lines even before the appearance of SR-mutations, indicating that mutations in other genes had also contributed to improvements in syntrophy. For instance, SR-mutations appeared before the 300th generation in HA2, UR1 and US1 but were not detected until the 500^th^ generation in HE3 and the 1000^th^ generation in HA3, HR2, UA3 and UE3 and not observed at all in HS3. DVU2894, a sigma-54 dependent regulator, was mutated in HA2 very early and later co-existed with SR-mutations. On the other hand, in UE3 mutations in DVU2894 appeared at the same time as SR-mutations, suggesting that multiple independent evolutionary trajectories could have led to improvements in syntrophy across the different lines.

In each evolution line, at least 8 (HR2) and up to 13 (HA3, UA3, and UE3) out of 24 high G-score mutations were selected in *Dv*, while *Mm* accumulated mutations in at least 5 (HA2, HR2 and, UA3) and up to 10 (HE3) out of 16 high G-score genes. The high degree of parallelism suggested that some populations could have converged to similar evolutionary trajectories due to epistatic interactions, in which case some mutations would frequently occur in the same order – because selection of one mutation allows for the next to be beneficial – or the alternative, that some pairs never occur together because of negative sign-epistasis. We interrogated the temporal order in which high G-score mutations were selected and the combinations in which they co-existed in each evolution line to uncover evidence for epistatic interactions in improving obligate syntrophy. Indeed, missense mutations in DsrC (DVU2776) were temporally correlated with the appearance of loss of function mutations in one of two sigma 54 type regulators (DVU2894, DVU2394) in lines HA2, UR1, UE3, and UA3. In rare instances, we also observed that some high G-score mutations co-occurred across evolution lines, e.g., DVU1283 (GalU) was mutated in two U- and one H-line and always co-existed with a mutation in DVU2394.

More commonly, the combinations in which high G-score genes accumulated mutations varied across multiple lines; in fact, no two lines possessed identical combination of high G-score gene mutations (**Fig 3A & B**). Many high frequency mutations were also uniquely present or absent in different lines (**Fig 3C and D**). For instance, in UR1 no mutations in DVU0799 were selected at any point through 1000 generations, even though this highest G-score gene in *Dv* was mutated in all of the other lines. Clonal interference (Maddamsetti et al., 2015) could also have limited parallel evolution in both *Dv* and *Mm* populations of UR1. In this line, mutations in DVU1214 and DVU2894 never achieved complete fixation, possibly due to competition with more beneficial mutations in DVU0013 and DVU2394. Similarly, mutations in MMP1362, MMP0335 and MMP1303 in the *Mm* population of UR1 appear to be outcompeted by mutations in MMP1718 and MMP0166.

**Figure 3.**
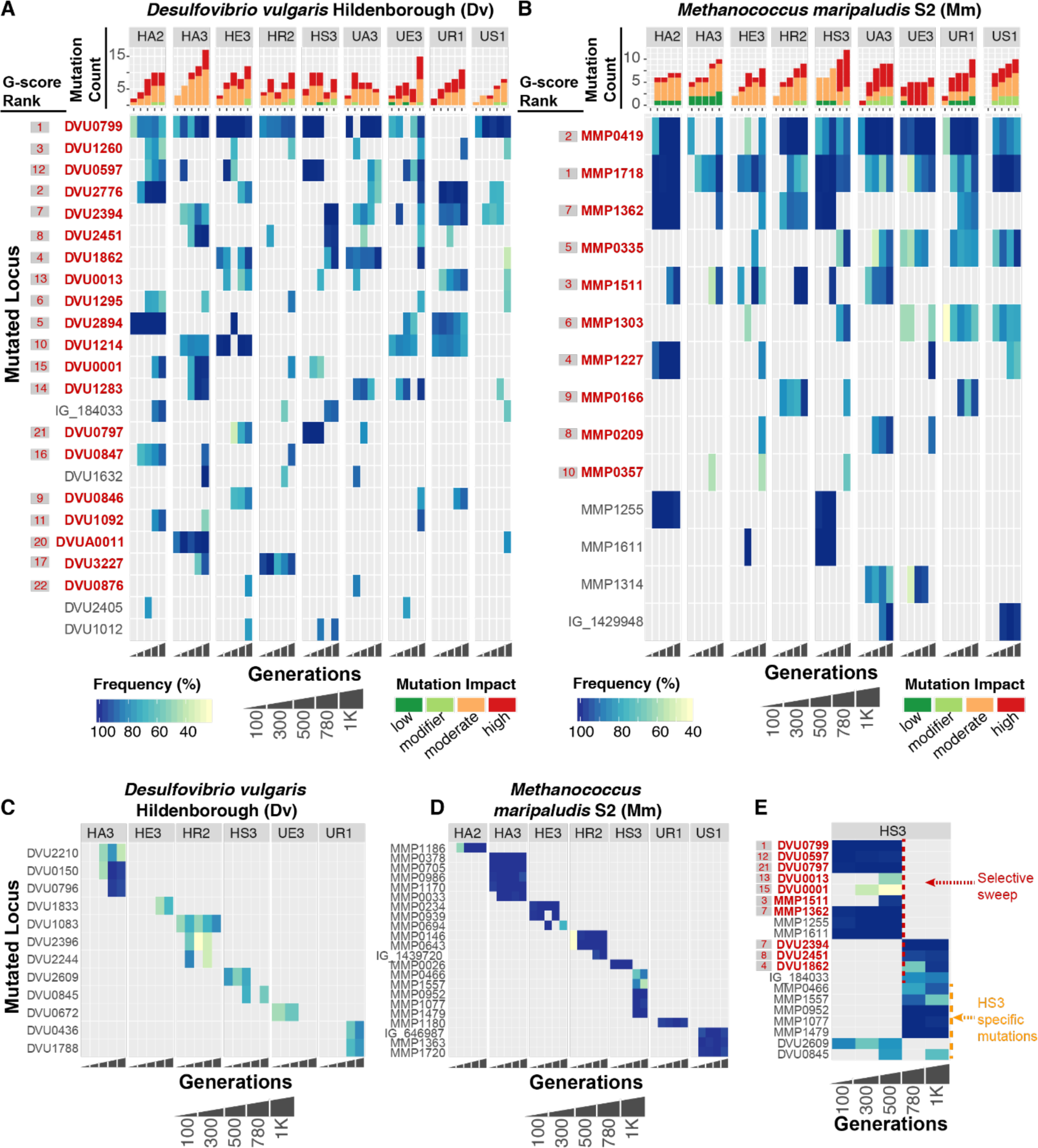
Frequency and time of appearance of mutations through 1K generations of laboratory evolution lines of *Dv* and *Mm* cocultures. The heat maps display frequency of mutations in genes (rows) in *Dv* **(A)** and Mm **(B)** in each evolution line, ordered from early to later generations (horizontal axis). High G-score genes are shown in red font and their G-score rank is shown to the left in gray shaded box, also in red font. Bar plots above heat maps indicate total number of mutations in each generation and the color indicates impact of mutation. Use “Frequency”, “Generations”, and “Mutation impact” key below the heat maps for interpretation. Mutations that were unique to each evolution line is shown in **(C)** and **(D)** for *Dv* and *Mm*, respectively. **(E)** The heat map illustrates a selective sweep across both organisms in line HS3.

The temporal order in which high G-score mutations accumulated within each line was also unique. Mutations in the same high G-score genes appeared at different times (e.g., whereas mutations in SR gene DVU0847 was first observed in the 300^th^ generation of HA2, they appeared much later in HR2 and HA3) (**Fig 3A**). Similar patterns of timing of appearance and co-occurrence of mutations were observed in *Mm* (**Fig 3B**). While mutations in MMP0419, MMP1718, MMP1227, and MMP1255 appeared within 100 generations of HA2, in the HA3 line most of these genes did not accumulate mutations until the 1000^th^ generation and no mutations were detected in MMP1227 and MMP1255. Similarly, all H-lines accumulated mutations in MMP1362 (DNA-directed RNA polymerase subunit beta), while only UR1 had this mutation among all U-lines. Conversely, all U-lines accumulated mutations in MMP0335 (hypothetical protein) across three different locations, but among H-lines, only HS3 had a mutation in this gene that was at the same location as in 2 U-lines (US1 and UE3).

We also discovered evidence for temporally nested fixations, wherein prior to fixation of a mutation from an earlier generation, another mutation selected in a later generation gradually increased in frequency towards fixation (e.g., DVU0799, DVU0001 and DVU1283 in HA3, MMP1718 and MMP0335 in UA3). Moreover, there were many cases of simultaneous fixation of mutations in multiple genes (e.g., DVU0799 and DVU1214 in HE3; MMP0378, MMP0705, MMP0986, and MMP1170 in HA2) suggesting that hitchhiking may be common (Maddamsetti et al., 2015, Lang et al., 2013). However, given that samples were only sequenced every 250 generations, we cannot rule out the possibility of each mutation sweeping separately during that time interval. These observations lead us to conclude that mutations that were commonly selected may simply have additive effects on fitness, arising at different times in different populations because of chance (i.e., they became available for selection at different times in different populations).

The longitudinal analysis revealed a cross-species selection event that resulted in the replacement of the dominant clones of both species with new clones containing different mutations. Between generations 500 and 780 in coculture HS3, the dominant Dv (harboring high G-score mutations DVU0799, DVU0597, DVU0797) and Mm (harboring dominant mutations in MMP1255, MMP1611, MMP1362, and MMP1511) clones disappeared (**Fig 3E**). At the same time a new Dv clone with mutations in DVU2394, DVU2451, DVU1862, and intergenic region IG_184033, and a new Mm clone with mutations in MMP0952, MMP1077, and MMP1479 were selected (**Fig 3E**). One explanation for this phenomenon is that rare clones in both Dv and Mm coincidentally each acquired beneficial mutations allowing them to outcompete dominant clones in the same 250 generation interval of evolution. Another possibility is that selection of a new dominant clone in one species changed the selection environment for the other, allowing its rare clone to take over. Which species might have started this process is unclear because there are no samples available in the 250 generations during which the swap occurred. However, information about the new mutations, their functions, and parallel evolution may provide hypotheses. The novel mutations in DVU2394 (a sigma54-dependent transcriptional regulator) and DVU2451 (a lactate permease) co-occurred in at least four lines including HS3 (the other three lines were HA3, UE2 and UE3) and appeared individually in only two other lines, suggesting that the two genes might be beneficial and also functionally coupled in the context of promoting syntrophy. Notably, we demonstrate later through single cell analysis that mutations in DVU2394 occurred subsequent to mutations in DVU2451, but exclusively in the SR-lineage within UE3. Interestingly SR-mutations were never selected through 1000 generations in HS3 line. Conversely, while mutations in DVU2394, DVU2451 and DVU1862, all co-occurred in UE3, they did not sweep through the population, underscoring how improvements to syntrophic interactions occurred through multiple distinct trajectories in terms of the order and combinations of mutation selection. In other words, this cross-species selective sweep occurred only in HS3, suggesting one of several features unique to this line was responsible, including simultaneous selection of mutations in DVU2394, DVU2451 and DVU1862, the overall mutational landscape of HS3 between generations 500 and 780, or mutations unique to HS3. Interestingly, fixed mutations that were observed only in HS3 were in *Mm* (MMP0952, MMP1077, MMP1479) and their appearance coincided with the selective sweep between 500 and 780 generations. The most plausible hypothesis based on these observations is that the selective sweep occurred due to loss of function mutations in MMP1077, a putative phosphomannomutase, which re-directed monosaccharides towards synthesis of exopolysaccharides to promote intercellular interactions through clumping or flocculation (Johnson et al., 2005). Regardless of the mechanism, it is especially interesting that a new mutation(s) in *Mm* appears to have selectively swept high G-score mutations across both members of the two-organism community, strongly suggesting that the new *Mm* genotype conferred a fitness advantage to a specific lineage of genotypes in *Dv* that were in low abundance prior to the sweep.

### Characterization of evolutionary lineages and interspecies interactions in minimal assemblages at single cell resolution

We performed end-point dilutions (EPDs) from the 1K generation of one heterogeneous (HR2) and one uniform (UE3) line to generate from each line simplified sub-communities that represent the minimal set of genotypes that display growth phenotypes comparable to the 1K culture (see Methods). While two EPDs from each of the two lines represented the dominant SR-subpopulation of the 1K evolved line, we also recovered an SR+ subpopulation that co-existed within each line albeit at much lower abundance and below the detection limit of bulk mutation analysis of the parental culture (**Fig 4**). Finally, we isolated evolved clones of each organism by streaking EPDs on agar plates containing nalidixic acid and neomycin, taking advantage of chromosomally-integrated selection markers for these antibiotics in *Dv* and *Mm*, respectively. Altogether, we obtained 3 clonal isolates of *Dv* and *Mm* from each EPD, and re-sequenced the genomes of these isolates. The distribution of unique mutations within these SR+ and SR-EPDs and 3 clonal isolates from each EPD added evidence for co-existence of distinct lineages of one or both organisms within each evolved line. Logically, all high frequency mutations in an asexual population must be linked on the same genetic background. As expected, all 15 high frequency mutations detected in the 1K generation of UE3 were present only in sub-communities with the SR-mutations (EPD-03 and EPD-10). By contrast, at least 11 mutated loci (10 genic and 1 intergenic) in the SR+ sub-community (EPD-09) were not detected in the SR-sub-communities or in 1K bulk sequencing of UE3, demonstrating that the EPD-09 assemblage was made up of rare *Dv* lineages (**Fig 4A** and Supplementary Table S1). Strikingly, both *Dv* and *Mm* lineages in the SR+ assemblage of HR2 were distinct from lineages in the SR-EPDs, and below detection limit in 1K bulk sequencing (**Fig 4B**, Supplementary Fig 1 and Supplementary Table S1). Thus, the isolation through dilution of genotypically distinct subpopulations of *Dv* and *Mm* having the parental growth phenotype was suggestive of the emergence of multiple interactions among specific evolved genotypes of the two organisms during their syntrophic evolution.

**Figure 4.**
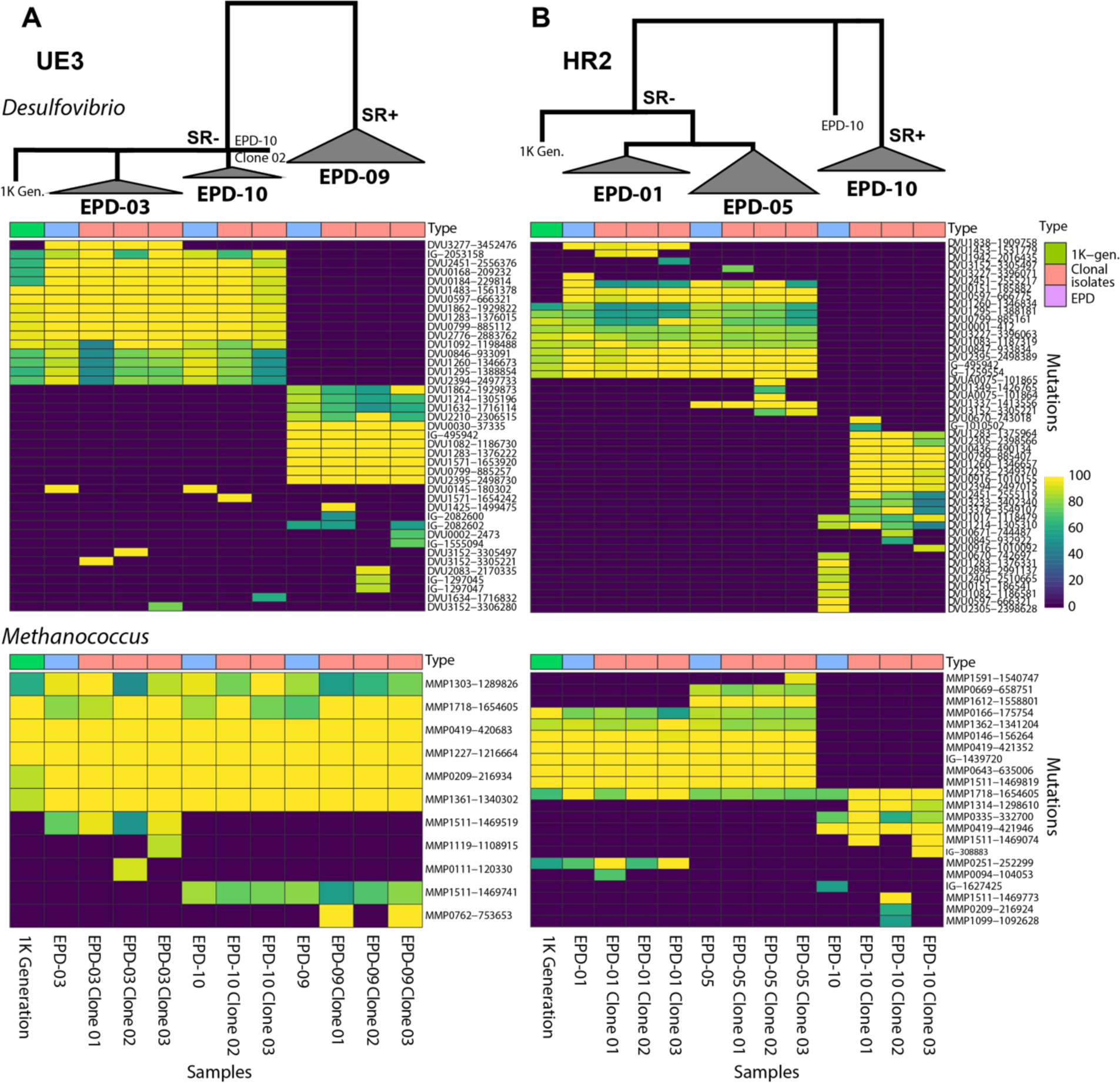
Genotype mapping of 1K generation, EPDs and clonal isolates for two evolution lines. Heatmap displays frequency for each mutation (rows) in UE3 **(A)** and HR2 **(B)** across 1K generation, EPDs, and clonal isolates (columns). The upper panel shows genotype map of *Dv* and the lower panel for *Mm*. The hierarchical tree indicates a simplified lineage map of mutations in *Dv* within each evolution line, with SR phenotypes indicated.

We further investigated evidence for interactions among specific evolved genotypes using single cell sequencing of SR- (EPD-03) and SR+ (EPD-09) assemblages from UE3. We sorted, amplified and re-sequenced the genomes of single cells of *Dv* (94 from EPD-03, and 94 from EPD-09) and *Mm* (87 from EPD-03, and 72 from EPD-09) to reconstruct lineages of both organisms within each EPD ((Thompson et al., 2017) and Methods, Supplementary Table S2). Altogether, 5,659,149 (EPD-03:4,596,604 and EPD-09: 6,721,694) and 321,310 (EPD-03: 240,853 and EPD-09: 401,767) reads mapped to the genomes of *Dv* and *Mm*, representing ∼225x and ∼29x coverage, respectively. Using stringent cut-offs (fold coverage ≥ 8, number of cells with mutation ≥ 2, frequency ≥ 80%) and consensus mutation calling using varscan (Koboldt et al., 2012), GATK (DePristo et al., 2011) and Samtools (Li, 2011), we identified across single cells of *Dv* 16 of 17 and 3 of 12 mutations detected in bulk sequencing of EPD-03 and EPD-09, respectively. Similarly, we identified across *Mm* single cells 7 of 7 and 6 of 7 mutations from bulk EPD-03 and EPD-09, respectively. Altogether, 70 mutations were shared across single cells of EPD-03 and EPD-09 (37 in *Dv* and 33 in *Mm*), and 11 EPD-specific mutations (1 in *Dv* and 10 in *Mm*) were not detected in bulk sequencing of EPDs, most likely because they were below the 20% frequency threshold of detection.

Using a mutation lineage inference algorithm SCITE (Jahn et al., 2016) and cross-referencing with longitudinal sequencing data from 5 generations (100, 300, 500, 780, 1000), bulk sequencing of EPDs, single cell sequencing, and sequencing of clonal isolates, we reconstructed the lineage and timeline of mutations that shaped the evolution of syntrophy in SR- and SR+ communities within UE3 (see Methods) (**Fig 5** and Supplementary Figures 2-3). As expected, the two EPDs shared a core lineage of events that included sequential accumulation of high G-score mutations in the early stages of evolution in both organisms. While the *Mm* lineages across EPDs had few differences, lineages of *Dv* were strikingly different across the SR- and SR+ communities. Of the total 11 high G-score *Dv* genes in the 1K generation of UE3, just three were observed in both EPDs. Strikingly, the three high G-score genes DVU1862, DVU2394, and DVU0799 had mutations in different locations in the two EPDs. High G-score genes that were only observed in EPD-03 were DVU2451, DVU1260 (outer membrane protein), and DVU1092 (Na-dependent symporter protein, and those unique to EPD-09 were DVU2395, DVU2210, and DVU1214. In addition, SR-mutations in DVU0846 and DVU1295 were unique to EPD-03, appearing after 780 generations, and were present across single cells and all clonal isolates. The SR-mutations in the EPD-03 lineage were followed by selection of mutations in at least six regulators, and complex radiating branches with many co-existing sub-clones, suggesting that loss of SR in the EPD-03 line might have promoted the selection of mutations in regulatory genes. Altogether, the observation that dominant lineages were excluded in the minimal community assemblages of EPD-09, demonstrates co-existence of distinct high abundance (SR-) and low abundance (SR+) lineages within the same evolved population (**Fig 4, Fig 5**, and Supplementary Figures 1-3). A surprising observation is that the SR+ clone that remained in the population subsequent to the evolution of SR-was not simply the dominant clone without the SR-mutation. Instead, it was a rare genotype with different mutations from the dominant population.

**Figure 5.**
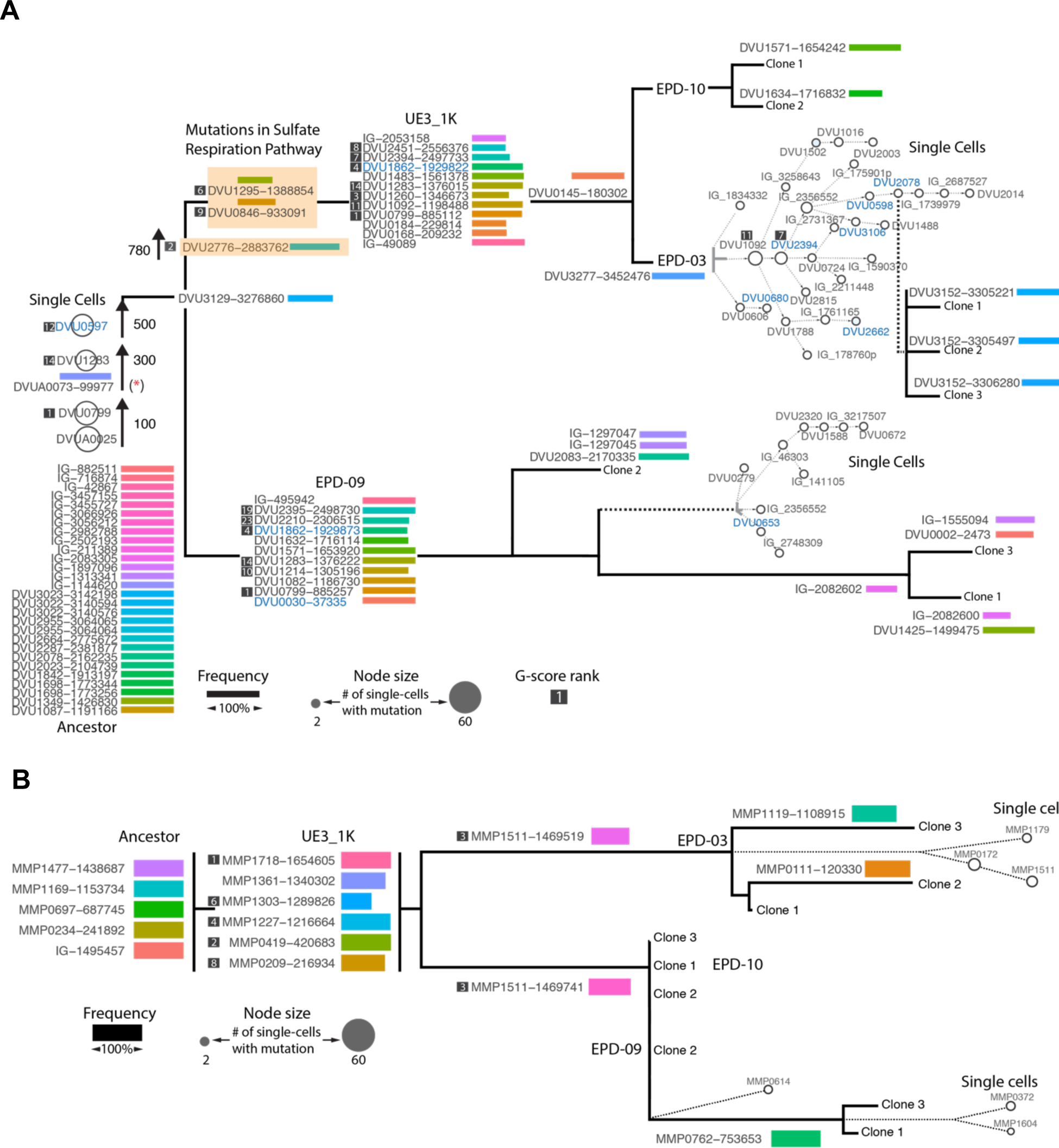
Lineage map of mutational events deciphered through sequencing of up to 96 single cells of **(A)** *Dv* and **(B)** *Mm* from EPD-03 and EPD-09, cross-referenced with longitudinal bulk sequencing of UE3, EPDs and sequencing of clonal isolates. Temporal ordering of mutations in the trunk is based on their order of appearance in longitudinal sequencing data across generations. Unique mutations within each lineage is shown together with frequency (length of bars). The single cell lineage tree for each EPD was constructed using the algorithm SCITE and shown in the context of the parent EPD and linked to clonal isolates. (See Supplementary Figures 2-3 for details). Mutation names for regulatory or signal transduction genes are colored in blue and SR-related genes are indicated with an orange shaded box. * indicates mutation in a plasmid gene that was not detected in single cells potentially due to loss of plasmid.

### Investigation of cooperativity and synergistic interspecies interactions

Given the possibility that interactions among specific genotypes of *Dv* and *Mm* had emerged during the evolution of syntrophy, we performed a density dilution assay to investigate evidence for improved cooperativity among microbial community assemblages of the two EPDs (Dai et al., 2012, Sanchez and Gore, 2013). Briefly, we generated an anaerobic dilution series of both EPD and ancestor cell lines in 96-well plates and experimentally determined growth rate, carrying capacity and a threshold dilution (i.e., minimal cell density) that supported syntrophic population growth (See Methods, Supplementary Figure 4). The density dilution assay revealed that both EPDs could initiate growth at significantly lower cell density relative to the ancestral coculture. EPD-03 initiated growth at a 1.5-fold lower cell density with faster growth rate and lower carrying capacity relative to EPD-09, explaining how the minimal assemblages represented by the two EPDs co-existed in vastly different proportions in UE3 (>80% EPD-03 vs, <1% EPD-09, **Fig 6A**). Collectively these data make a compelling case for the emergence of increased cooperativity among *Dv* and *Mm* lineages during the laboratory evolution of syntrophy.

**Figure 6.**
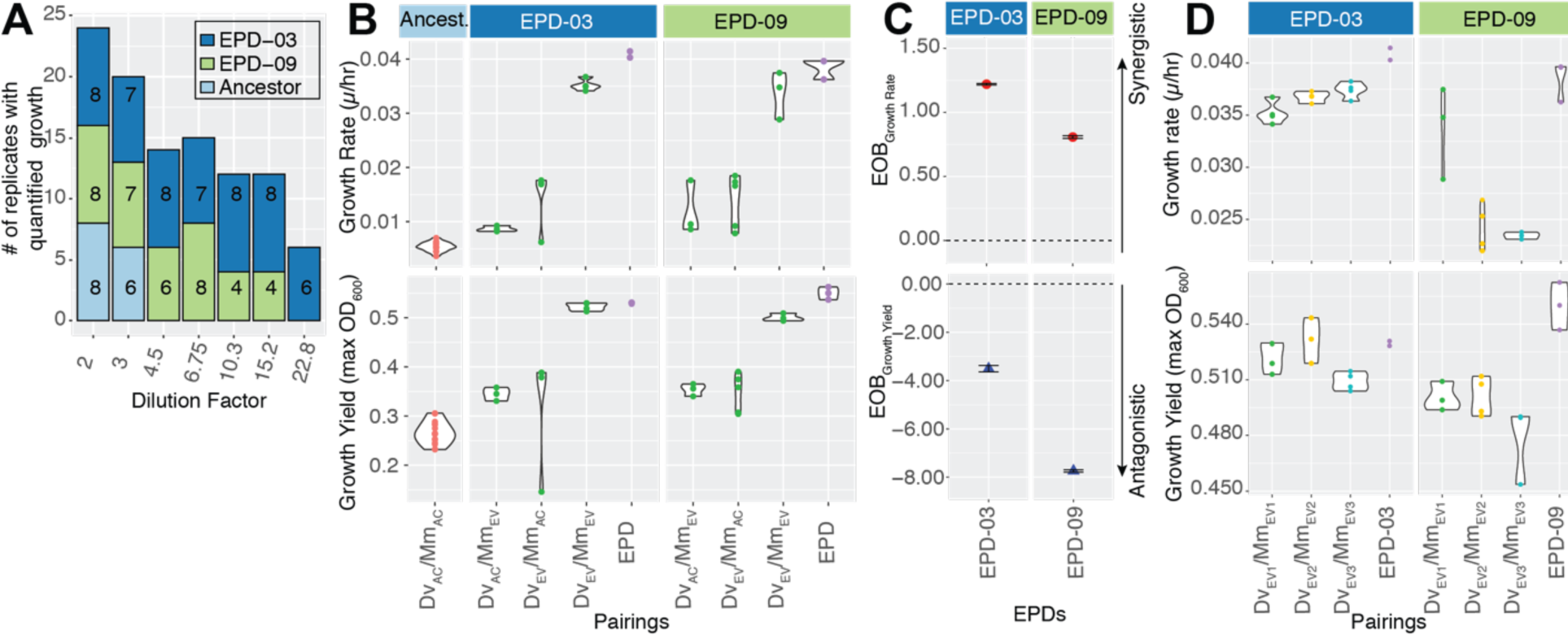
Growth rate, yield and cooperativity of EPDs, and clonal isolate pairings. **(A)** A stacked barplot showing the number of replicates exhibiting growth for each EPD and the ancestral cocultures across a dilution series. **(B)** Growth rate and carrying capacity of pairings of ancestral and evolved clonal isolates of *Dv* and *Mm* from EPD-03 and EPD-09. **(C)** Excess-Over-Bliss analysis for estimating synergistic and antagonistic interactions of *Dv*/*Mm* clonal isolate pairings. (**D**) Growth rate and yield for 3 evolved *Dv*/*Mm* pairings from each EPD.

To further investigate whether increased cooperativity had emerged through the evolution of synergistic interspecies interactions, we characterized individual and combined contributions of the two evolved partners towards improved growth rates and yields. Based on growth characteristics of pairings of evolved isolates with each other (*Dv*_EV_ x *Mm*_EV_) and their ancestral counterparts (*Dv*_Ac_ x *Mm*_Ev_ and *Dv*_Ev_ x *Mm*_Ac_), we determined that each evolved clonal isolate of *Mm* and *Dv* had contributed individually to significant improvement in growth rate and yield, relative to the ancestral pairing (*Dv*_Ac_ x *Mm*_Ac_) **(Fig 6B** and Supplementary Table S4). The improvements were maximal, and comparable to growth characteristics of the parental EPD, when both partners in the interacting pair were evolved clonal isolates (*Dv*_Ev_ x *Mm*_Ev_). This result demonstrated unequivocally that increased cooperativity had emerged from synergistic interactions between the evolutionary changes in both species within each EPD, with proportional antagonistic effect on growth yield (Novak et al., 2006) (**Fig 6C**). The higher growth rate of EPD-03 and higher carrying capacity of EPD-09 (both relative to the other EPD) gives mechanistic insight into co-existence of SR- and SR+ sub-communities as *r*- and *K* strategists, respectively (**Fig 6D**, Supplementary Figure 4). Notably, the few mutations that differentiate genotypes of each clonal isolate appear to manifest in variation in growth rate and yield, demonstrating that productivity of *Dv*_*EV*_ x *Mm*_*EV*_ interactions are genotype-specific, even within the same EPD (**Fig 6D**, Supplementary Figure 5, Supplementary Table 3).

## DISCUSSION

We sought to understand the evolutionary trajectories that increase the productivity of interspecies interactions of *Dv* with *Mm* in an obligate syntrophic association, while retaining a small subpopulation that can respire sulfate. To do so, we combined a broad survey of all the mutations accumulated over the first 1000 generations of 9 independently-evolved communities with an in-depth study of the genotypic structure of one community down to the single-cell level. These data showed a high level of parallelism across communities in the genes that acquired mutations despite considerable variance across populations in their evolutionary trajectories. A detailed view of one community revealed the perseverance and evolution of a rare lineage that maintained its ability to respire sulfate while the rest of the population did not. Growth experiments with clones and subpopulations demonstrated that the SR+ and SR-*Dv* subpopulations both cooperate more efficiently with corresponding evolved *Mm* partners, allowing them to grow at lower starting densities than the ancestors. The collective action of clones within each subpopulation has a synergistic effect on population growth rate and an antagonistic effect on yield. Finally, the different growth dynamics of SR- and SR+ evolved communities explained how the two communities co-exist in vastly different proportions as *r*- and *K*-strategists.

The evolutionary trajectory of a microbial population depends on the order in which mutations occur (chance), and the relative effects of the pool of mutations on fitness (selection) (Travisano et al., 1995). If the effects of each beneficial mutation are constant, meaning they do not vary in the presence of other polymorphisms or species, then all populations would eventually acquire the same mutations, even if they occur and are, therefore, selected in a different order. However, the effect of an allele on fitness may depend on epistasis, where the effect of an allele changes depending on alleles at other loci in the same genome (Wolf et al., 2000). In this case, the order in which mutations occur in different populations could affect their overall trajectories. For example, a beneficial mutation that nullifies the effects of other beneficial mutations could force a population down a different evolutionary trajectory from those that acquire the nullifying mutation last or never at all. This relationship between fitness and the possible combinations of genetic variants is called an adaptive landscape, and has been the subject of intense research (Orr, 2005, Wright, 1932, Weinreich et al., 2006, Flynn et al., 2013).

In the present work, we investigated evolutionary trajectories of not just one but two species that rely on one another for survival. One might expect the interaction to amplify the effects of chance if the adaptive landscape is affected by genetic changes in the partner population (coevolution; (Thompson, 1989, Hillesland, 2018)), and those partner genetic changes depend on chance. In an extreme case, this situation could send each population down completely different trajectories, with very little parallelism. However, that is not the result that was observed here. The discovery of high G-score mutations in both *Dv* and *Mm* made a compelling case that parallel evolution was a dominant driver of productive obligate syntrophy (**Fig 2**). Genes associated with parallel evolution are usually under strong selection (Bailey et al., 2017), and implicated as a major driver of evolution of bacteria (Woods et al., 2006), phages (Wichman et al., 1999) and microbial communities (Douglas et al., 2016). The genes that were subject to parallel evolution across multiple lines were associated with either loss of function (e.g., DVU1295, DVU0846, MMP0335 etc.), function modulation (e.g., DVU2776, DVU0597, MMP1227 etc.) or both types of mutations (e.g., DVU0799, DVU2394, MMP0419, etc.). To our knowledge, this is the first demonstration of a role for parallel evolution in driving mutualism across metabolically coupled species.

While the high number of G-score mutations suggests that parallel changes conferred fitness benefits across a range of adaptive mutation landscapes (Wichman et al., 1999), in some populations these high G-score mutations were selected in a different order, suggesting epistasis did not substantially constrain when high G-score mutations could be selected. Many differences between populations in mutation order could have occurred due to the chance occurrence of mutations at different times in different populations (Lenski et al., 1991). However, we cannot rule out the possibility that epistasis and evolutionary history caused some of the differences between populations (Blount et al., 2018, Elena et al., 1996). For example, it is possible that a mutation unique to HS3 precluded or significantly delayed erosion of SR in this coculture.

Another conclusion that could be drawn from the high degree of parallelism is that the interaction between *Dv* and *Mm* has little effect on their evolution, and thus each species is essentially evolving alone in a constant environment consisting of another species. However, there are a few reasons to believe this is not entirely correct, and that some evolutionary changes likely resulted from genetic interactions between specific evolved genotypes of *Dv* and *Mm*. The most striking example was observed in community HS3. In this community, it seems that one or more new mutations in *Mm* (or Dv) affected the fitness of the dominant *Dv* (or Mm) clone, causing it to decrease in frequency below the limits of detection, while a new clone arose. This two-population selective sweep demonstrates epistasis between genotypes of the two interacting species. In addition, growth rates and yields differed between some pairings of clones within a population, demonstrating variation in effectiveness of cooperation.

While most adaptive mutations rose to fixation, SR mutations did not show complete penetrance. In fact, previously we reported that SR+ populations were readily obtained from every evolved coculture even after 1,300 generations (Hillesland et al., 2014) and through characterization of EPDs and single cells we have re-confirmed the presence of rare SR+ subpopulations in most evolved populations. One explanation could be that SR+ cells persist because maintenance of sulfate-respiration machinery allows them to produce a costly but essential metabolite. Leaking of this metabolite could allow SR-cells to survive without paying the cost of production, allowing them to flourish as long as SR+ cells and the leaked resource do not become scarce. In other words, these SR+ cells might act as “helpers” for the “beneficiary” SR-cells as stated by Black Queen Hypothesis (Morris et al., 2012). High expression of SR genes even under syntrophic conditions (Walker et al., 2009) supports this hypothesis. However, a minimal assemblage that is entirely composed of SR-cells (EPD-03 of UE3 line) does better than assemblage of SR+ cells (EPD-09 of the same line) in cooperativity assays with no apparent growth defect indicating that SR+ cells do not play an essential role in supporting syntrophic growth of the population. In fact, the poor performance of EPD-09 relative to EPD-03 suggests that SR is too expensive to maintain and, therefore, undesirable during syntrophy. Moreover, individual SR-clonal isolates synergistically improved growth characteristics of cocultures upon pairing with evolved *Mm* further demonstrating that co-existence of SR+ and SR-populations cannot be explained solely by the BQH.

Alternatively, SR+ and SR-cells may be adapted to different niches that arise as a result of the seasonal changes in resources that recur in each transfer-cycle of the evolution experiment (Rozen and Lenski, 2000). Specifically, growth dynamics of the two EPDs [higher growth rate (*r*) of EPD-03 and higher carrying capacity (*K*) of EPD-09] suggest that the faster growing SR-lineages (*r*-strategists) can initiate growth at lower cell density and are, therefore, favored in early growth phase when resources are plentiful but fluctuating. The slower growing SR+ lineages (*K*-strategists) are favored in later stages of growth when the resources are limited but stable, and cell density is high (Wei and Zhang, 2019). Hence, these growth dynamics based on *r/K* tradeoffs might explain why SR+ populations are retained in the absence of sulfate. In the natural world, where sulfate availability varies over time, persistence of SR+ genotypes in the absence of sulfate may stabilize sulfate-reducing populations overall. In other words, it may be a bet-hedging strategy (similar to maintenance of subpopulations with COO hydrogenase polymorphisms (Großkopf et al., 2016, Beaumont et al., 2009) that might contribute to the success of *Dv* as a generalist that can conditionally switch between SR and syntrophy without the need for expensive gene regulatory changes (Turkarslan et al., 2017).

It was significant that each of the two EPDs segregated a subset of high G-score mutations into simplified assemblages but retained growth rate and carrying capacity of the parental evolved population. This result demonstrated that multiple independent evolutionary strategies can co-exist in the same population, albeit in vastly different proportions. Whether the distinct sets of *Dv* and *Mm* mutations within each EPD reflect coevolution will require additional experiments, including pairing with evolved populations from preceding generations (Hillesland, 2018). Notwithstanding that caveat, the ability of evolved isolates of *Dv* and *Mm* to synergistically improve growth characteristics lends credibility to the claim that complementary genetic changes (e.g., in transport, regulation, and motility) enhanced metabolic coupling and cross-feeding between the two interacting organisms, significantly increasing their cooperativity.

The nature of cooperation in this syntrophic mutualism is unclear. On the surface, it seems like the fitness of Dv and Mm would be aligned and exploitation unlikely (Marx, 2009, Estrela and Brown, 2013, Oliveira et al., 2014) because the production of hydrogen is a necessary byproduct of metabolism for Dv and the only energy source available for Mm. Efficient transfer of electrons through hydrogen is in the best interests of both species (Stolyar et al., 2007). However, evolution could hypothetically change this situation by altering mechanisms of electron transfer, or through the evolution of new dependencies that are costly (Harcombe et al., 2018, Hillesland, 2018). One high G-score mutation in Mm (MMP1511) could reflect the evolution of a new costly dependency. Alanine was earlier shown to be exchanged between the two interacting partners during syntrophic growth, likely at a cost to the producer (*Dv*) and of energetic advantage to *Mm* (Walker et al., 2012). Alanine production by *Dv* provides a mechanism to re-oxidize reduced internal cofactors during syntrophic growth, but at the cost of a high-energy phosphate bond. In turn, alanine taken up and converted to pyruvate and ammonia by *Mm* serves as both a carbon and a nitrogen source, alleviating complete dependency on energetically costly autotrophic growth with hydrogen. A cheater population, e.g., one with a loss of function mutation in MMP1511, might consume additional alanine through passive transport and therefore consume less hydrogen to maintain lactate consumption by *Dv*. Indeed, mutations in MMP1511 rose to fixation in six out of nine lines, and we cannot rule out if minor MMP1511 mutant populations also exist in low frequency in the other lines, including UE3.

The observation that interactions among some genotypes were more productive than other pairings suggests that the enhanced cooperativity of evolved communities could have occurred through the selection of complementary mutations across *Dv* and *Mm*, invoking the possibility of partner choice and partner fidelity feedback (Archetti et al., 2011). Furthermore, each EPD had significantly better growth characteristics than any of the pairings of their member clonal isolates, demonstrating the emergence of increased cooperativity from guilds or “collections of genotypes” of *Dv* and *Mm*. In conclusion, the multiscale dissection of independent laboratory evolution lines has demonstrated that selection of complementary mutations across *Dv* and *Mm* synergistically increased the cooperativity and productivity of syntrophic interactions within both SR- and SR+ communities, while supporting their co-existence in vastly different proportions as *r*- and *K*-strategists, respectively.

## ACKNOWLEDGMENTS

This material by ENIGMA-Ecosystems and Networks Integrated with Genes and Molecular Assemblies (http://enigma.lbl.gov), a Science Focus Area Program at Lawrence Berkeley National Laboratory is based upon work supported by the U.S. Department of Energy, Office of Science, Office of Biological & Environmental Research under contract number DE-AC02-05CH11231. In addition, sequencing of ancestral cocultures and early generation samples was supported by the National Science Foundation under Grant No. DEB-1453205 and DEB-1257525 to KLH. The pipeline for sequencing data analysis was developed using funds from the National Institute of Health under grant number R01AI141953 to NSB. We would like to thank Nicholas Elliott for his help with growth analysis, Joseph Hellerstein, and Adrian Lopez Garcia de Lomana for discussion on data analysis.

## AUTHOR CONTRIBUTIONS

Conceptualization and Methodology, S.T., N.S., N.S.B., D.A.S. and K.L.H; Investigation, S.T., N.S., A.W.T., C.E.A., J.J.V., J.W., K.A.H., J.H., Y.F., L.W., and Y.M.S.; Formal Analysis, S.T., N.S., and N.S.B.; Data Curation, S.T., N.S., K.L.H., K.A.H., and N.S.B; Writing – Original Draft, S.T., N.S.B., K.L.H., and D.A.S.; Visualization, S.T., N.S.B.; Supervision, N.S.B., K.L.H., and D.A.S.; Resources and Funding Acquisition, J.Z., N.S.B., K.L.H., and D.A.S.

## METHODS

### Strains and Culture Conditions

All the strains, culture conditions and the setup of the laboratory evolution experiment were the same as described before (Hillesland and Stahl, 2010, Hillesland et al., 2014). Briefly, two clones of *Desulfovibrio (Dv)* and *Methanococcus (Mm)* were paired to setup 24 ancestral cultures in coculture medium A (CCMA) (Stolyar et al., 2007) under anaerobic conditions (80% N_2_:20% CO_2_ headspace) in Balch tubes. Cocultures were propagated weekly into a fresh media through 100-fold dilutions and incubated either upright without shaking or in a horizontal position with constant shaking at 300 rpm. Laboratory evolution experiment was continued for 152 weeks and populations were archived as frozen glycerol stocks after generations 100, 300, 500, 780, and 1000 generations. Biomass collection was done as described before (Hillesland et al., 2014).

### Sequencing of Evolved Cocultures

DNA sequencing was performed for 13 of 22 evolved cocultures after 1000-generation and 9 evolved cocultures were sequenced at 100, 300, 500 and 780-generations. In addition, three End-Point-Dilutions from UE3 (EPD-03, EPD-09 and EPD-10) and HR2 (EPD-01, EPD-05 and EPD-10) cocultures evolved for 1000 generations and 3 clones of *Dv* and *Mm* from each of these EPDs were sequenced. For each sample, DNA was extracted with Epicentre Masterpure Kit (Epicentre Catalog number: MC85200). Sample and sequencing library preparation was done by using the Nextera DNA library preparation kit (Illumina) according to the manufacturer’s instructions. DNA sequencing was performed in an Illumina Hiseq (generations 100, 300, 500, and 780) with 100 bp paired end sequencing or in an Illumina MiSeq sequencing instrument in the paired-end mode producing 2×250 bp long reads as described before (Hillesland et al., 2014).

### Identification of Mutations in Evolved Cocultures

Mutations accumulated in populations were determined by using a custom sequence alignment and variant calling pipeline (https://github.com/sturkarslan/evolution-of-syntrophy). This pipeline included quality control and trimming of the raw sequencing reads in fastq format by using Trim Galore software (http://www.bioinformatics.babraham.ac.uk/projects/trim_galore). The alignment of the quality trimmed sequences to reference *D. vulgaris* (Genbank assembly: GCA_000195755.1.30) and *M. maripaludis* (Genbank assembly: GCA_000011585.1) genomes and subsequent processing steps before calling the variants was done by following The Genome Analysis Toolkit (GATK) (DePristo et al., 2011) best practices. Briefly, reads were first aligned to the reference genome using Burrows-Wheeler Alignment Tool (bwa) (Li and Durbin, 2009) (version 0.7.17-r1188) in paired-end mode. The resulting alignment files in the SAM format were converted to BAM files, sorted and indexed by using Samtools version 1.9 (Li et al., 2009). BAM files were marked for duplicates using Picard Tools (http://broadinstitute.github.io/picard/) (version 1.139), and local realignment around indels was performed to identify the most consistent placement of reads relative to the indels. Variant calling was performed independently by using three different algorithms including GATK UnifiedGenotyper, Varscan (Koboldt et al., 2012) (version 2.3.9) and bcftools from Samtools package. The default parameters were used for UnifiedGenotyper, whereas for Varscan parameters were --min-coverage 8 --min-reads2 2 --min-avg-qual 30 and bcftools parameters were -vmO -s LOWQUAL -i’%QUAL>30. Variants identified by each caller were collated and filtered for variant frequency equal or greater than 20%. A variant was included in the analysis only if it is simultaneously called by at least two of the callers. The resulting variants were annotated using SnpEff tools (Cingolani et al., 2012) (version 4.3).

### Single Cell Sequencing

For single cell sequencing, EPD-03 and EPD-09 from UE3 evolved cocultures were grown to mid-log phase. Single cells of *Desulfovibrio* or *Methanococcus* were sorted into wells of a 96-well plate containing 3 µl of PBS and Buffer D2 from Repli-G single cell kit (Qiagen) by using Influx flow cytometer (BD). For each EPD, one plate for each of *Desulfovibrio* and *Methanococcus* was prepared. In order to lyse the cells, a freeze-and-thaw cycle was performed by first spinning the plates and freezing them at −20°C followed by thawing and re-spinning. Whole Genome Amplification (WGA) from single cells was performed by using REPLI-G Single Cell kit (Qiagen) according to manufacturer’s instructions. We screened single amplified genomes (SAGs) with 16S universal primers for *Desulfovibrio* or *Methanococcus* to identify percentage of wells that did not contain any amplified product due to missing cells or failed WGA reaction. Wells with confirmed amplification were further treated with AmpPure XP magnetic beads (Beckman-Coulter) to clean and purify SAGs. A subset of SAGs was also analyzed with Bioanalyzer to confirm the size of the amplified fragments. Concentration of the SAGs passing the quality controls were determined by using Quant-iT PicoGreen dsDNA assay kit (Thermofisher). Nextera XT Library preparation kit (Illumina) was used for sample and library preparation for sequencing. Sequencing was performed in HiSeq platform (Illumina) by using High-Output flow cell in 2×150 bp paired-end format. Sequence analysis including quality controls, trimming, alignment and variant calling was performed as described above.

### Single Cell Lineage Tree Building

Variants identified from single cells of *Desulfovibrio* or *Methanococcus* for EPD-03 and EPD-09 of 1000-generation evolved UE3 cocultures were converted into a binary mutation matrix where each row was a unique mutation and each column was a single cell. Values in the binary mutation matrix were either 1 (mutation was seen in that particular cell), 0 (mutation was not observed) or 3 (there wasn’t enough confident reads to assign the mutation). A variant was considered for the analysis only if its frequency was over 80% and was seen in at least two single cells. Mutation histories of single cells were determined by using SCITE algorithm (Jahn et al., 2016) with parameters -r 1 -l 900000 -fd 6.04e-5 -ad 0.21545 0.21545 -cc 1.299164e-05. SCITE used stochastic search to find the Maximum Likelihood tree of mutation histories in Newick format, which was converted to Cytoscape format for visualization purposes. This tree represents the predicted temporal order of the mutation events. Mutations were re-ordered by using information from the sequencing of the early generation cocultures if the order of the mutations couldn’t be determined from the single cell mutational profiles due to noisy and missing data. Mutation tree was further annotated with gene functions, type of mutations and status of the mutations in early generations, and clonal isolates.

### Calculation of G-scores

Based on the frequency of observed mutations (normalized to gene length and genome size) across 13 evolved lines, we calculated a G-score (“goodness-of-fit”) to assess if the observed parallel evolution rate was higher than background as described before (Tenaillon et al., 2016). Briefly, expected number of mutations (*E*_*i*_) for each gene in the genome was calculated as:

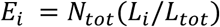

where *N*_*tot*_ is the total number of mutations, *L*_*i*_ is the length of the gene *i* and *L*_*tot*_ is the total length of the coding genome. G-score for each gene (*G*_*i*_) was calculated as:

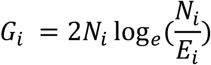

where *N*_*i*_ is the number of nonsynonymous mutations observed for gene *i* across all evolved lines. G-scores for all genes in the genome of each organism were summed up to get the “total observed G-statistic” (*G*_*obs*_). In order to get the “total expected G-statistic” (*G*_*exp*_), we simulated N_tot_ number of mutations randomly across the protein-coding genome and calculated the mean and standard deviation of G-statistic from all simulations. We compared the observed and expected G-statistics by calculating a Z-score as follows;

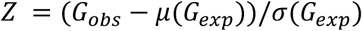

Where *μ*(*G*_*exp*_) and *σ* (*G*_*exp*_) are the mean and standard deviation of the G-statistics from 1000 simulations, respectively.

### Density dilution assay

Ancestor and EPD cocultures were revived anaerobically in 18×150-mm balch tubes (Chemglass Life Sciences: CLS420901) from freezer stocks through dilution into CCMA media to ensure syntrophic growth and prevent carryover of glycerol into the fresh growth medium. EPD batch cultures (10 mL) were grown in anaerobic conditions with 80%:20% N_2_:CO_2_ headspace at 30°C without shaking. Growth of the cultures were monitored using a spectrophotometer (Spectronic 200: Fisher Scientific) to measure the optical density at 600 nm (OD600), measurements were typically taken twice a day until the cocultures reached stationary phase (∼0.7 for EPD-03 and EPD-09). The cultures were kept in stationary-phase for approximately 20 hours in order to ensure similar growth phases before dilution into 96-well plates (Thomas Scientific: 1154Q44) for the density dilution assay. Stationary-phase Ancestor and EPD cultures were all diluted to the same starting optical density (as measured by the 96-well plate reader: BioTek EPOCH2T), so that all cultures in column 12 of each plate have the same starting optical density. Thereafter, each column between 11 and 2 received a volume of cells that would equate to a 1.5-fold dilution of the previous column and starting with column 11. The 1^st^ column of each 96-well plate contained only media as a control to identify potential contamination. All dilutions of cocultures into plates for the density dilution assay were done inside a Coy Anaerobic Chamber with an approximate atmospheric ratio of 95%:5% N_2_:H_2_. Plates were sealed with optically clear strong adhesive PCR films (115×100mm, Thomas Scientific: 4ti-0500/8), and the edges of these seals were coated twice in clear acrylic to further inhibit potential gas diffusion into the wells. Following inoculation, sealed plates were incubated at 30°C within the anaerobic chamber. Plates were removed from the anaerobic chamber twice per day to take growth measurements in the BioTek plate reader. Plates in the plate reader were shaken linearly for 5 seconds prior to OD600 measurements at 30°C. During the transfer of plates from anaerobic chamber to plate reader, they were insulated between two 6-well plates filled with H_2_O that were also incubated at 30°C in order to maintain constant temperatures during plate transport from anaerobic chamber to plate reader. Density dilution assays were carried out for approximately ∼4-5 days. A moving average with a window of two was applied to ODs for each density dilution assay to smooth the timeseries data. In order to establish a baseline for growth, a threshold was calculated for each EPD and ancestor strain based on the minimum carrying capacity from the first two dilutions (n=16). If cells from a well did not achieve an OD of that minimal threshold or higher, those cells were considered as not grown.

### Clonal isolate pairings and measurement of Growth Rate and Yield

Isolates of *Desulfovibrio* from EPD-03 or 09 of line UE3 were revived anaerobically using 10 ml of CCMA containing 10 mM:7.5 mM sodium lactate:sodium sulfite or 30 mM:20 mM sodium lactate:sodium sulfate, respectively, in balch tubes flushed with 80%:20% N_2_:CO_2_. Isolates of *Methanococcus* were revived anaerobically using 5 ml of CCMA containing 10 mM sodium acetate in balch tubes pressurized to 30 psig with 80%:20% H_2_:CO_2_. After the second transfer of revived isolates on their respective media, 0.1-0.2 ml of stationary phase cultures were combined in 20 ml of CCMA containing 30 mM sodium lactate in balch tubes flushed with 80%:20% N_2_:CO_2_. After the second transfer, cocultures were stored as freezer stocks for future growth analysis. Cocultures for growth analysis were revived anaerobically using 20 ml of CCMA containing 30 mM sodium lactate in balch tubes flushed with 80%:20% N_2_:CO_2_. Cultures were incubated at 37°C and shaken horizontally at 300 rpm. Optical densities (OD_600nm_) were monitored to assess growth and growth parameters were estimated using the fitting package grofit (Kahm et al., 2010).

### Excess over Bliss analysis for measuring synergy

We adapted the Bliss Independence model (Borisy et al., 2003) to predict if accumulated mutations in evolved *Dv* and *Mm* partners have an additive effect on growth rate and yield of their clonal isolate pairings. The experimentally measured fractional growth rate and yield for *Dv* (*f*_*Dv*_) and *Mm* (*f*_*Mm*_) was determined by pairing their evolved clonal isolates with ancestral clones of their respective partners. Then, the expected fractional effect on growth rate and yield *f*_*DvMm*_, induced by the combined effect of evolved isolates was calculated as:

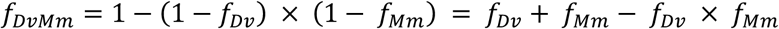

Excess over Bliss (EOB) was determined by computing the difference in fractional improvement of growth rate or yield induced by combination, *f* _*z*_ and the expected fractional inhibition, *f*_*DvMm*_

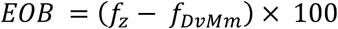

A clonal isolate pair combination for which EOB ≈ 0 has an additive behavior, whereas a pair with positive or negative EOB values has synergistic or antagonistic behavior, respectively. Error bars were computed by propagating the standard deviation of fractional effects.

### Data and Code availability

Bulk and Single cell sequencing data used in this study and associated biosample meta-data information can be obtained through the NCBI Bioproject database (https://www.ncbi.nlm.nih.gov/bioproject) with accession number PRJNA248017. Custom R and Python codes used for sequence analysis, variant calling, data analysis and figure preparations are available on GitHub (https://github.com/sturkarslan/evolution-of-syntrophy). Annotated mutations within the context of other functional and regulatory genome information can be explored through Syntrophy Portal (http://networks.systemsbiology.net/syntrophy/)

## SUPPLEMENTARY FIGURES

**Supplementary Figure 1.**
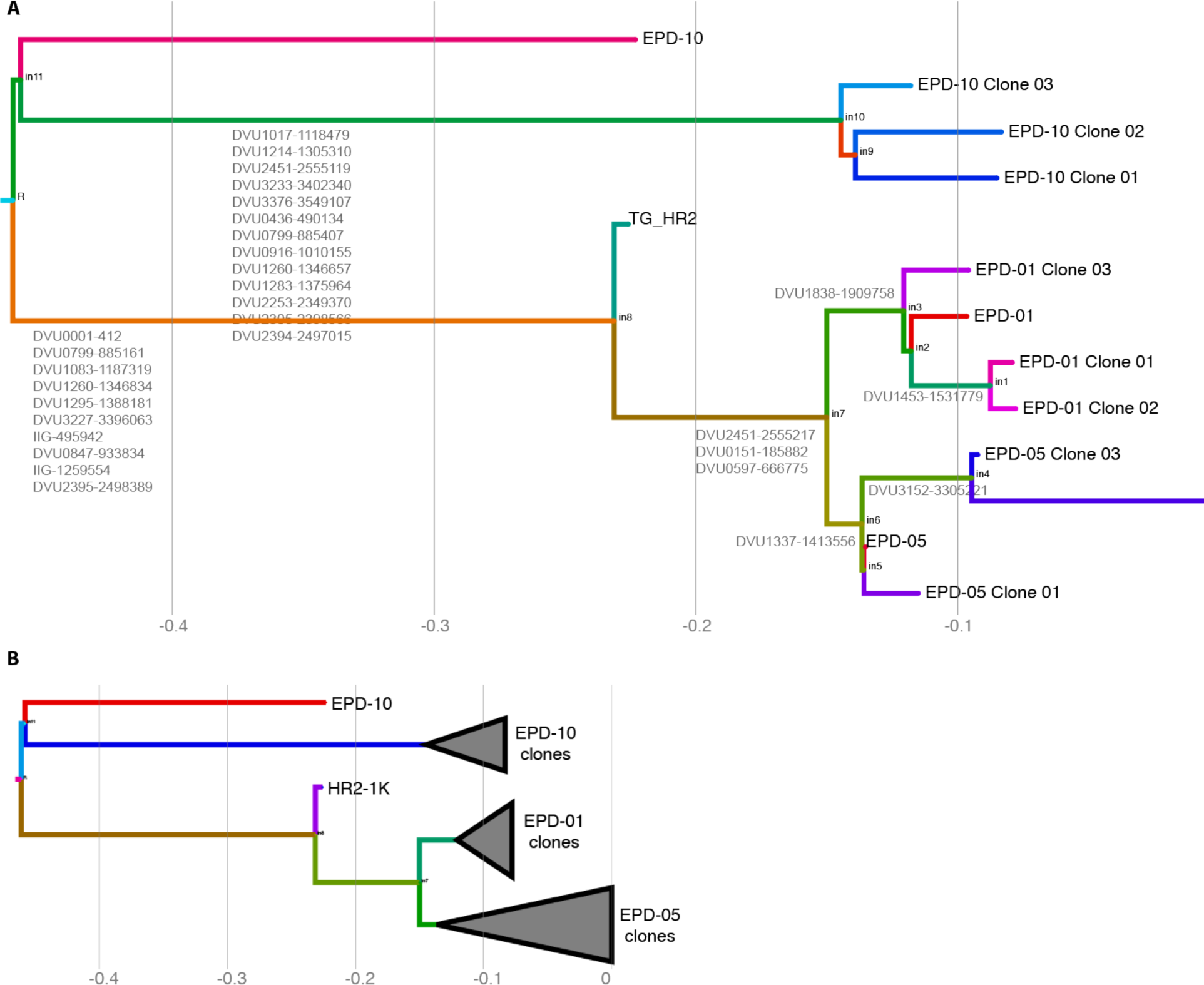
HR2 lineage tree with mutations mapped to branches. All mutation data (except early generations) for HR2 including 1K generation, EPDs and Clonal isolates were used to discover lineage relationships between EPDs by using siFit algorithm (Zafar et al., 2017) and tree was plotted by using IcyTree (Vaughan, 2017). **A)** Detailed tree **B)** Simplified tree with lower tree branches collapsed.

**Supplementary Figure 2.**
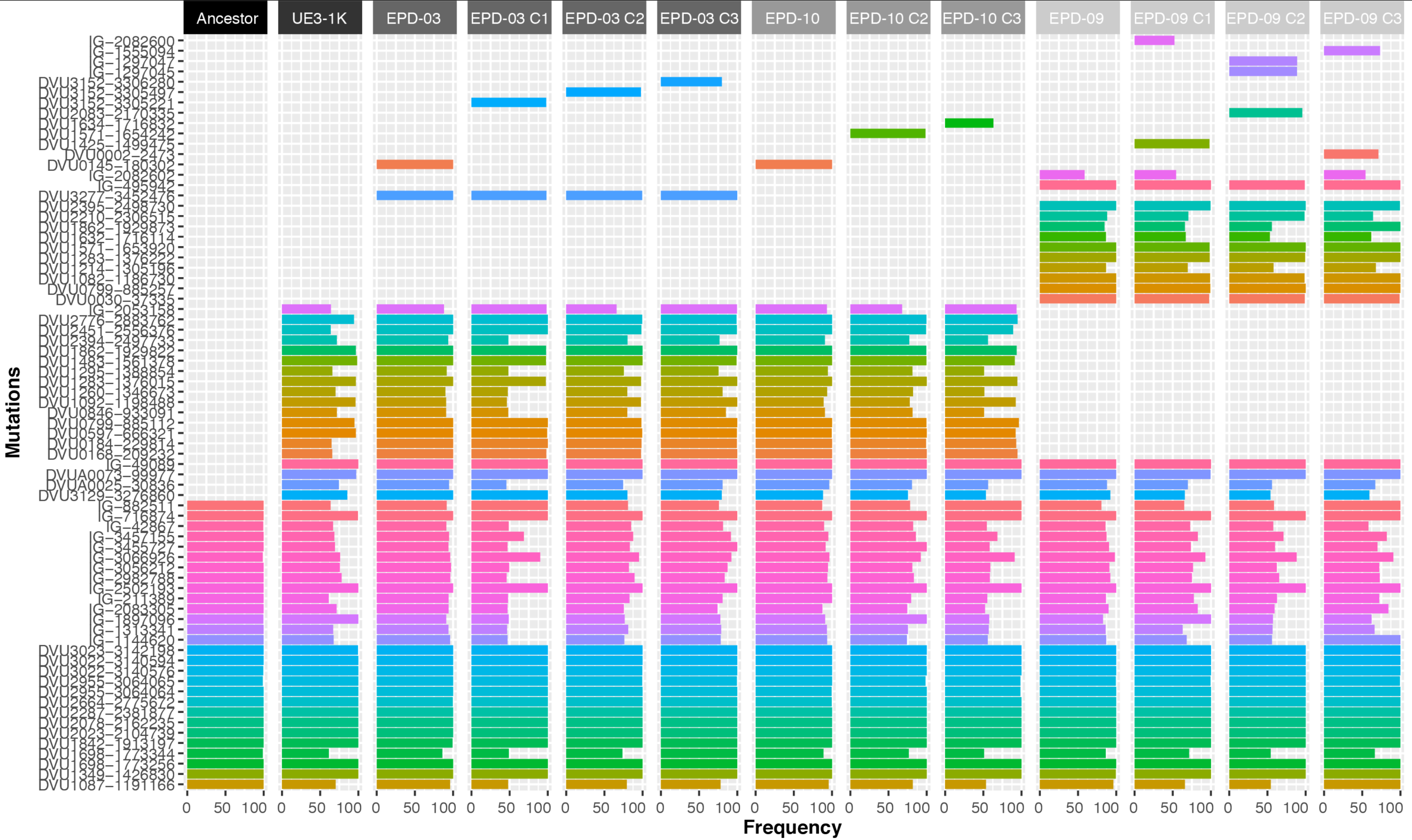
Comparison of *Dv* mutations identified in ancestral cocultures and accumulated in UE3 line across, 1K generation, EPDs and Clonal isolates. The length of bar plots denotes frequency for each mutation (rows) while color indicates different mutations.

**Supplementary Figure 3.**
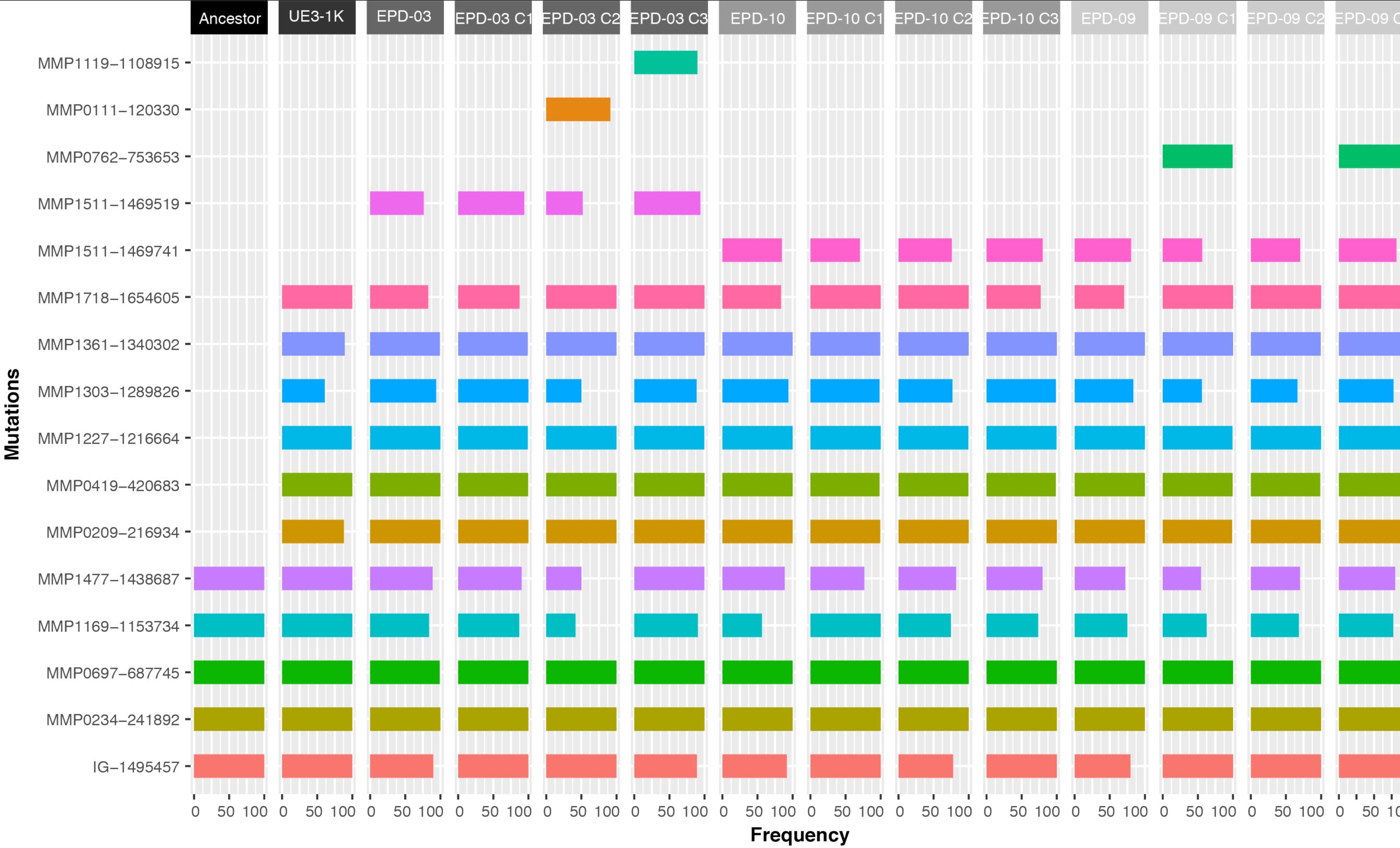
Comparison of *Mm* mutations identified in ancestral cocultures and accumulated in UE3 line across, 1K generation, EPDs and Clonal isolates. The length of bar plots denotes frequency for each mutation (rows) while color indicates different mutations.

**Supplementary Figure 4.**
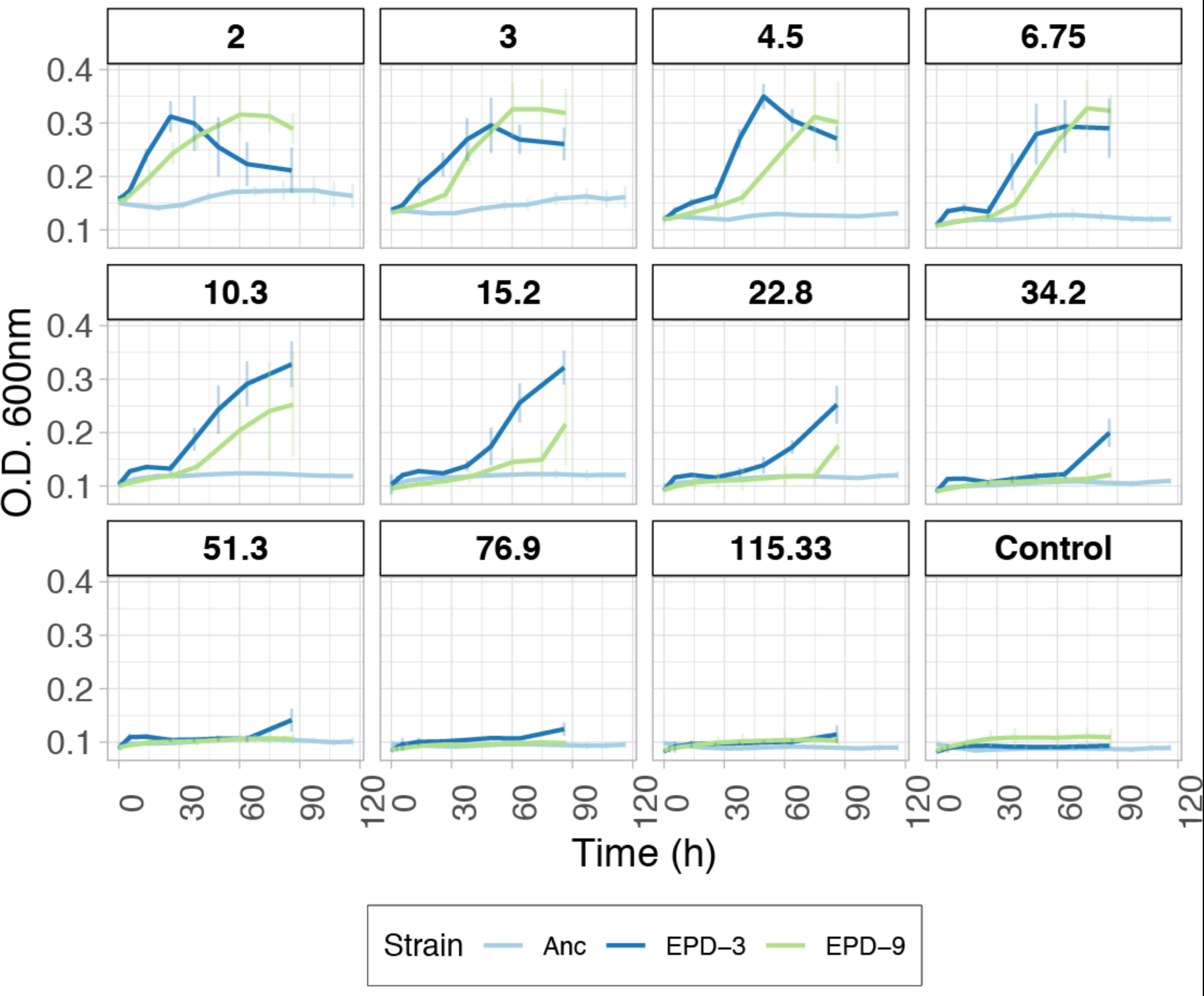
Density dilution assay growth curves to quantify cooperativity. Using 1.5 serial dilutions in 96-well plates, the Ancestor (Anc), EPD-03, and EPD-09 were grown anaerobically in plate reader and growth was followed by OD measurements as described in the Methods. Dilution factors are indicated as panel titles.

**Supplementary Figure 5.**
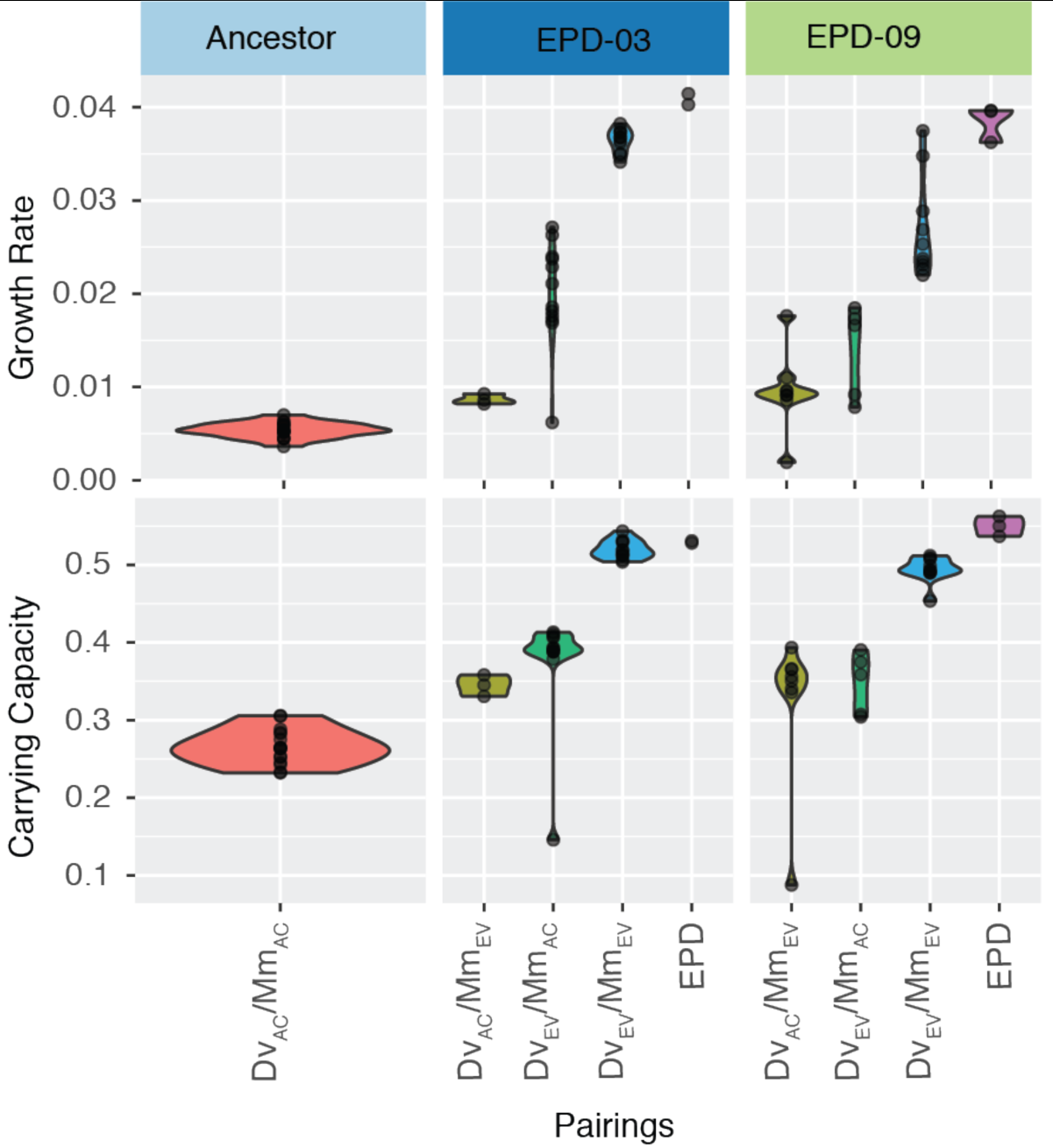
Growth rate, yield and cooperativity of EPDs, and clonal isolate pairings.

